# Mapping fetal myeloid differentiation in airway samples from premature neonates with single-cell profiling

**DOI:** 10.1101/2022.07.08.499395

**Authors:** Holly Welfley, Ranjit Kylat, Nahla Zaghloul, Marilyn Halonen, Fernando D. Martinez, Mohamed Ahmed, Darren A. Cusanovich

## Abstract

Single-cell genomic technologies hold great potential to advance our understanding of development and disease. A major limitation lies in isolating intact cells from primary tissues for profiling. Sampling methods compatible with current clinical interventions could enable longitudinal studies, the enrollment of large cohorts, and even the development of novel diagnostics. To explore single-cell RNA-seq (scRNA-seq) profiling of the cell types present at birth in the airway lumen of extremely premature (<28 weeks gestation) neonates, we isolated cells from endotracheal aspirates collected from intubated neonates within the first hour after birth. We generated data on 10 subjects, providing a rich view of airway luminal biology at a critical developmental period. Our results show that cells present in the airways of premature neonates primarily represent a continuum of myeloid differentiation, including fetal monocytes (25% of all cells), intermediate myeloid populations (48% of cells), and macrophages (2.6% of cells). To our knowledge, this is the first single-cell transcriptomic characterization of human monocytes in the neonatal airway isolated within an hour of birth. Applying trajectory analysis to the premature neonate myeloid populations, we identified two trajectories consistent with the developmental stages of interstitial and alveolar macrophages, as well as a third trajectory presenting a potential alternative pathway bridging these terminal macrophage states. While the three trajectories share many dynamic genes (5,451), they also have distinct transcriptional changes (259 alveolar-specific genes, 666 interstitial-specific genes, and 285 bridging-specific genes). Overall, our results define high quality single-cell data from cells isolated within the so-called “golden hour of birth” in extremely premature neonate airways representing complex lung biology and can be utilized in studies of human development and disease.

## Introduction

Single-cell RNA sequencing (scRNA-seq) technologies are transforming our understanding of human development, complex disease etiology, and disease progression. This is in large part due to the ability of single-cell techniques to discriminate distinct cell populations within heterogeneous tissues [1] and allow for the inference of cellular differentiation and cellular response trajectories within an individual cell type. In fact, scRNA-seq techniques have already led to the discovery of two novel cell types in the lung [2–4]. Despite these advances, a major challenge of single-cell profiling remains the ability to obtain representative suspensions of cells (or nuclei) of the tissue of interest from human patient samples [5–7]. Provided access to appropriate tissues, scRNA-seq studies could help to inform clinicians on appropriate therapeutic interventions and provide insights into normal and diseased cellular function over time. Therefore, an important factor in the ultimate utility of single-cell approaches is the identification of samples that can be collected during routine clinical practice and serve as sentinels for their source tissues.

In the case of respiratory diseases, one tool has been the profiling of endotracheal aspirates to reflect biology of the airway. Although limited to individuals who have been intubated, the aspirates allow a window into airway development and disease by profiling cells of the airway lumen [8]. Aspirates open the possibility of observing cells from the airways of neonates within minutes of birth (for children experiencing respiratory distress), as well as adults intubated for diseases like pneumonia. Transcriptional profiles of endotracheal aspirates have been shown to be a useful tool in characterizing respiratory diseases in adults, including ventilator-associated pneumonia [9], as well as distinguishing between COVID-19-associated acute respiratory distress syndrome (ARDS) and ARDS arising from other etiologies using scRNA-seq [10]. In premature infants, endotracheal aspirates have been used to identify biomarkers of subsequent development of bronchopulmonary dysplasia (BPD) from protein levels in aspirate fluid [11].

Nonetheless, studies of tracheal aspirates with scRNA-seq remain limited. This is particularly true for aspirates of premature neonates. To date, we are aware of only one study applying scRNA-seq to aspirates from two premature neonates [12]. Due to a limited number of cells profiled and an emphasis on epithelial cell types, the cellular annotations and transcriptional profiles were not fully resolved (with some cell clusters left unannotated by the authors). Although animal models, organoid models, air-liquid interface cell culture models, etc. have been used to characterize lung development at the cellular and molecular level [13, 14], and human tracheal aspirate samples have been investigated with flow cytometry and bulk RNA-seq [15], a complete scRNA-seq reference for the cell types present in human airways throughout gestation and after birth remains to be developed. For example, the source and timing of development of alveolar macrophages has been of interest in the field. Previous work in human subjects, using histology or flow cytometry on samples from stillborn neonates, noted that alveolar macrophages were not typically observed in the lungs of neonates until 24 to 48 hours after birth, leaving open the question of how these important immune cells derive from fetal precursors in humans [16, 17]. While this process is well worked out in mice [18], we are not aware of any previous report of the transcriptional profile of definitive fetal monocytes (presumed alveolar macrophage progenitors) or the trajectories of their differentiation in human airways. Two studies have presented results of single-cell RNA-seq analysis of fetal lung tissue. One study examined lung tissue from subjects that were at 11.5 and 18.5 weeks gestational age [19] and identified primarily mesenchymal cells with some epithelial, lymphoid, and myeloid cells becoming more apparent at the later time point. A second study profiled lung tissue from a range of ages spanning approximately 12 to 20 weeks gestational age [20], and identified seemingly more differentiated stromal and epithelial cells, as well as smaller, less characterized clusters of lymphoid and myeloid cells (the latter of which the authors subsequently classified as perivascular macrophages). Studies have also characterized lung tissue samples from subjects 29-31 weeks gestational age via scRNA-seq. However, these were collected 1-4 days after birth in subjects with major congenital anomalies [21]. Therefore, holes remain in our understanding of the transcriptional regulation of the developing human airways.

Studying the aspirates of extremely premature neonates (<28 weeks gestational age) in particular presents a unique opportunity to observe the state of airway cells at a critical stage in development. The periviable period of birth (the youngest gestational age at which neonates are able to survive birth) is 22 to 26 weeks gestational age [22], although other factors are more predictive of viability than gestational age alone [23]. This developmental window overlaps the transition from the canalicular stage of lung development (16-26 weeks gestational age) to the saccular stage (the final prenatal stage of lung development, 24-38 weeks gestational age) [24]. Given the importance of this time window in development and viability, further characterization is warranted. Furthermore, the perinatal period of lung development for extremely premature neonates is a critical factor in the potential for developing life-long disorders such as BPD, pulmonary hypertension, neurological complications, increased risks of infections, asthma, and a need for ongoing breathing support [25]. Advancements in perinatal care, such as ventilation and administration of surfactant, have drastically improved patient outcomes. However, the major clinical determinate of disease is failure to thrive which occurs after the onset of symptoms [26], highlighting the need for earlier intervention, which might be facilitated by better understanding of the developmental and disease processes occurring in the airways of these children.

To define the cells present in the airways in this critical developmental window (23-28 weeks gestational age), we performed scRNA-seq on tracheal aspirates collected upon initial intubation collected from a cohort of extremely premature neonates. Beyond the potential for enabling diagnostic and intervention development, exploring these samples provides a baseline understanding of the developmental stage of luminal cells during the canalicular/saccular transition. Importantly, by integrating our results with previously published lung single-cell profiles in adults, we were able to compare the cell types observed in the airway at different stages of development and to expand our perspective on the developmental state of the myeloid compartment of the airway at birth. While the difference in sampling technique (tracheal aspirate vs. lung tissue) and health status of the samples (premature neonates in need of intubation vs. adults without lung disease) undoubtedly influences which cellular populations may be surveyed, we found the integration of the two data sets useful in framing meaningful differences between them.

## Results

### Endotracheal aspirates from extremely premature neonates are enriched for myeloid populations

To generate profiles of airway cells at birth for extremely premature neonates, we collected endotracheal aspirate samples within an hour of birth upon being intubated by clinical directive from premature neonates with a gestational age of 23-28 weeks (gestational birth weight < 1250 gram). Samples were collected and subsequently cryopreserved to be processed in batches. While overall sample viability and cell yield were relatively low, we were able to generate data on 6,803 cells across 10 subjects after quality filtering (See Methods for details, Table S1, Fig. S1). Visualizing data from all 10 individuals together in a Uniform Manifold Approximation and Projection (UMAP) plot (Fig. 1A), we identified 11 clusters of cells (Fig. S2). To annotate cell types, we first used several label transfer approaches - including the label transfer algorithm from Seurat [27] with lung references from both Travaglini et al. [28] and the healthy subjects from Adams et al. 2020 [29], as well as SingleR [30], which uses a different algorithm and a different set of transcriptional reference profiles (Fig. S2, S3, Table S2). Subsequently we defined differentially expressed marker genes using a likelihood-ratio testing framework applied to ‘pseudobulk’ gene expression estimates for each individual for each cell cluster to identify cell type marker genes [31]. This analysis identified 438-2,550 marker genes for each cluster (Fig. 1B,C, Table S3). Finally, we considered the correlation of pseudobulk gene expression estimates for each cluster to determine how distinct the expression patterns of related clusters were (Fig. S2D). Considering all three analyses (label transfer, differential expression, and correlation analysis results), we assigned 9 cell types to the 11 clusters, with two pairs of clusters being merged primarily because of similarities in cell type annotations and gene expression profiles). Across our cohort, we identified a large population of myeloid cells (6 of the 9 cell types, including classical monocytes, macrophages, and neutrophils) and two smaller clusters of epithelial cells, which we annotated as basal and ciliated cells.

**Figure 1:**
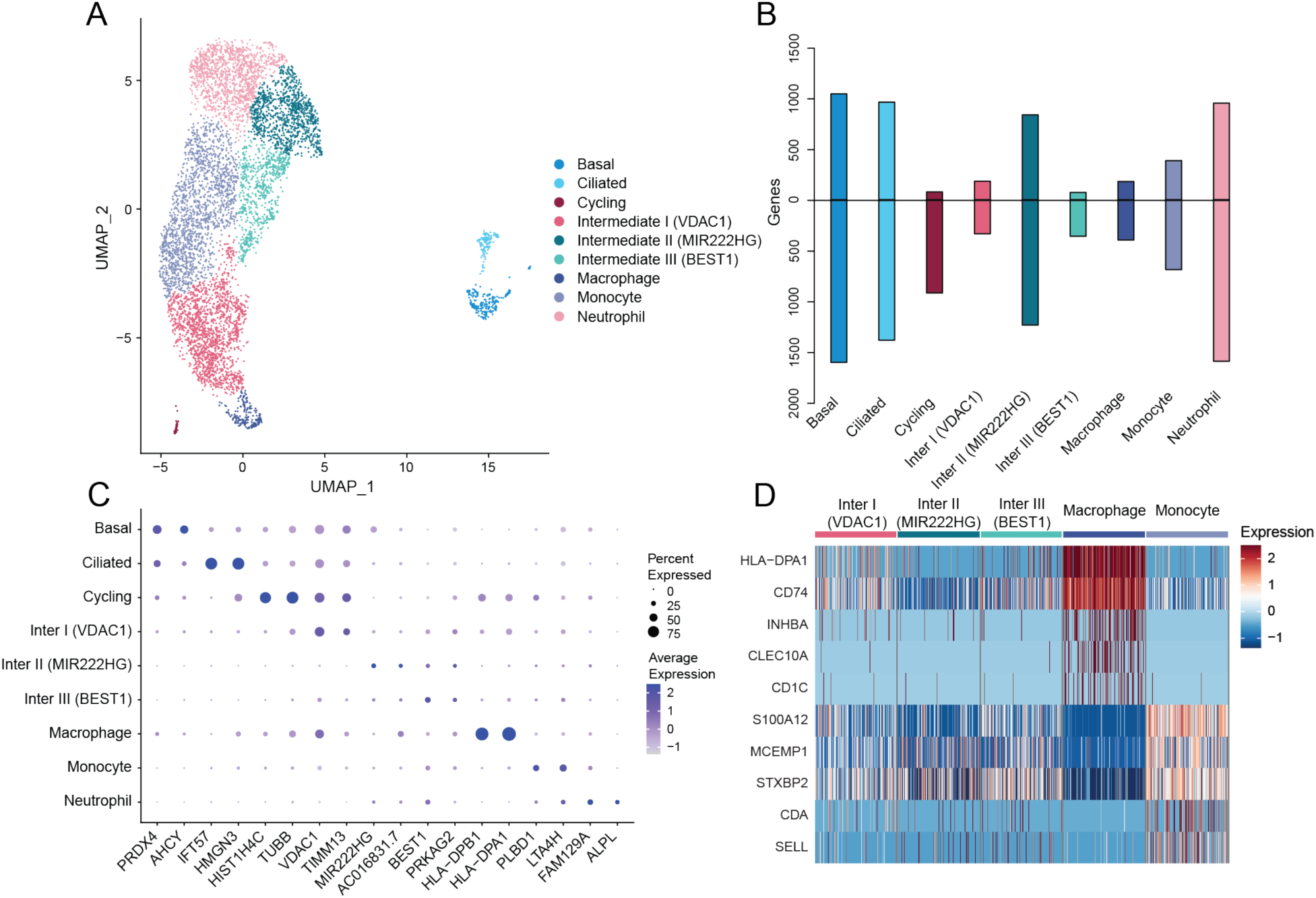
Cell type annotation of premature neonate endotracheal aspirate samples. (A) UMAP visualization of cell types identified across 10 extremely premature neonate aspirate samples. (B) Bar plot showing the number of significant (adjusted p-value < 0.05) differentially expressed genes for each annotated cell type. Bars above the line indicate genes with a Log2(fold-change) > 0 (“upregulated”) for that cluster relative to all other cell types. Bars below the line indicate genes with a Log2(fold-change) < 0 (“downregulated”) for that cluster. (C) Dot plot highlighting the top two most significant marker genes for each cell type. Each row represents an individual cell type. Each column is a marker gene. The diameter of the dot indicates the fraction of cells in that cell type expressing that gene. The color of the dot indicates the average expression level of that gene in the cell type. (D) Heatmap of the top five most significantly upregulated genes in monocytes and macrophages relative to each other (in cases where multiple genes from the same family were in the top 5, only the most significant member of the family is included). Each row represents a gene and each column represents an individual cell. Cells from each cell type were randomly subsampled down to 177 cells for visualization (to match the number of cells in the smallest cluster, macrophages).

**Figure 2:**
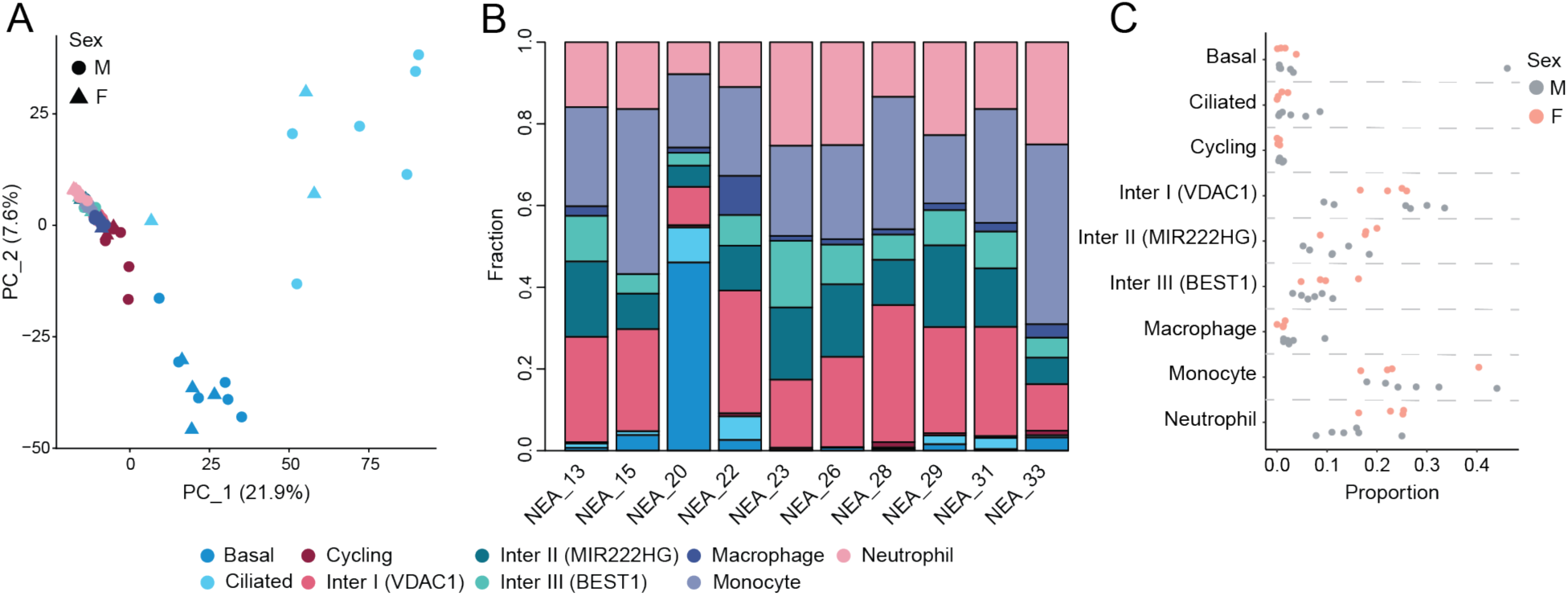
Cell type distribution and expression by patient and sex. (A) PCA plot of the pseudobulk profiles of each defined cell type (indicated by point color) per subject (shape indicates the sex of the subject). (B) Barplot of the cell type composition within each patient. (C) Stripchart showing the proportion of cells assigned to each cell type per individual stratified by sex.

**Figure 3:**
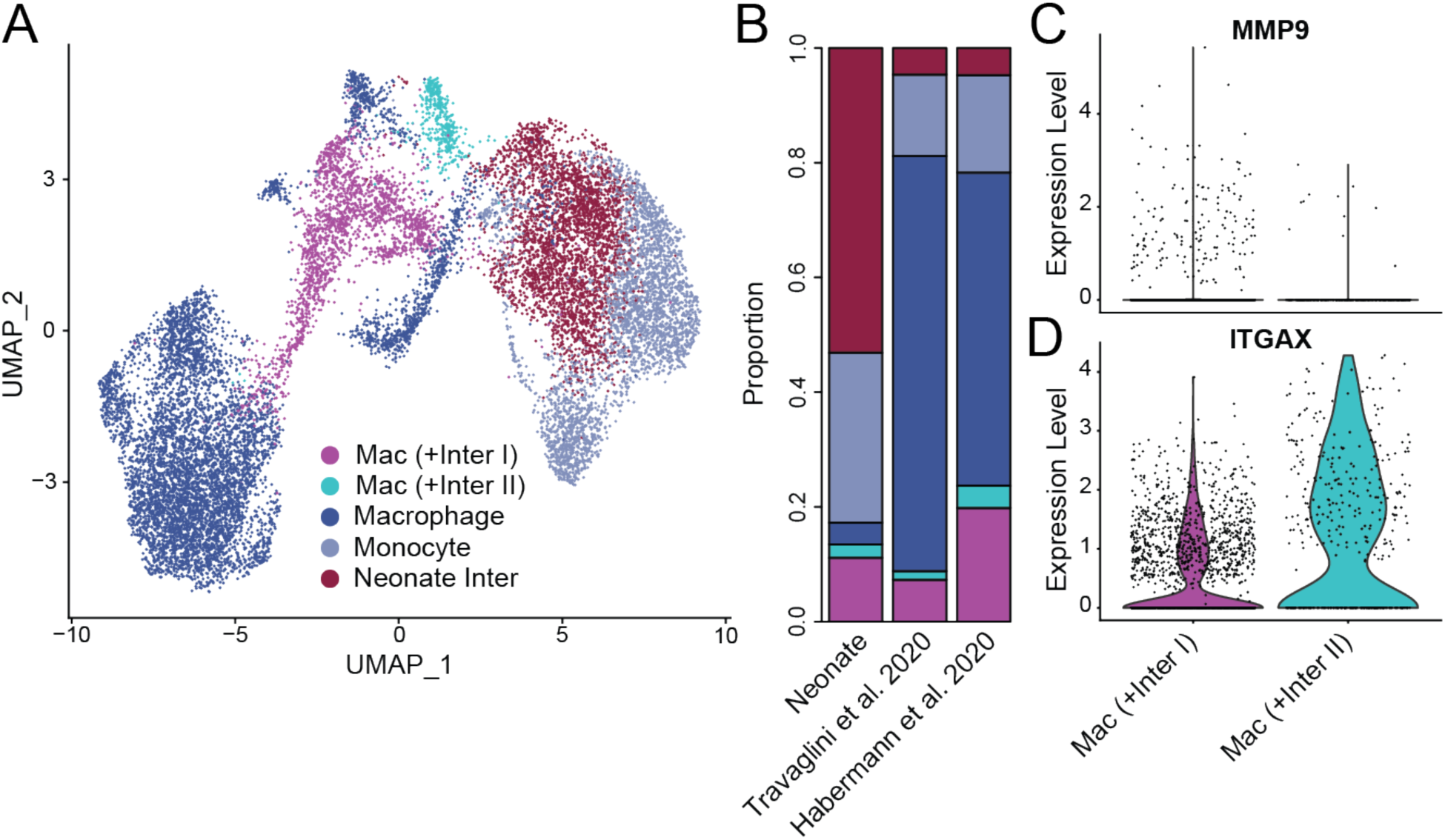
Integration with adult lung myeloid cells. (A) UMAP visualization of monocytes, intermediate myeloid cells, and macrophages integrated across premature neonate aspirate samples and two healthy adult lung tissue published datasets. (B) Barplot of the cell composition from each contributing dataset, Travaglini et al. [28], Habermann et al. [37]. (C-D) Violin plots of interstitial macrophage marker *MMP9* (C) and alveolar macrophage marker *ITGAX* (D) expression across macrophage populations co-clustering with premature neonate intermediate cells.

The myeloid compartment contained a large cluster of monocytes and a small cluster of differentiated macrophages. In addition, four myeloid cell clusters that had reduced expression of canonical markers for progenitor cells (i.e. monocytes), such as *S100A8*, *S100A9*, and *FCN1*, or for more differentiated types (i.e. macrophages and dendritic cells), including *C1QB*, *MRC1,* and *MSR1*, and the label transfer algorithms assigned these cells with less confidence or inconsistent annotations (Table S2, Fig. S4A-F). Therefore, we sought to better define these clusters by additional analyses. Given that the samples were initially stored on ice and then cryopreserved before single-cell profiling, we were somewhat surprised to find that one of these clusters (Cluster 1, Fig. S2C) displayed lower overall RNA complexity and was positive for previously determined markers of neutrophils (Fig. S4G-I), such as *FCGR3B* (adjusted p-value = 2.9x10^-10^; [32]), *CXCR2* (adjusted p-value = 1.9x10^-08^; [33]), and *CMTM2* (adjusted p-value = 2.0x10^-11^; [34]). For the remaining three myeloid clusters, we performed differential expression analysis between monocytes and macrophages and examined the expression pattern across the three clusters of genes upregulated in monocytes or macrophages, as well as considering the overall transcriptional correlation of these three clusters with monocytes and macrophages (Table S4, Fig S5). We found that none of these unannotated myeloid clusters consistently expressed the top genes differentiating monocytes and macrophages (Fig. 1D), and the correlation structure of their transcriptional profiles placed them intermediate to monocytes and macrophages via unsupervised hierarchical clustering. Therefore, we annotated these clusters as “Intermediate” and differentiated them based on the most significantly differentially expressed gene for each cluster: “Intermediate I (VDAC1)”, “Intermediate II (MIR222HG)”, and “Intermediate III (BEST1)” (Fig 1C).

### Sex differences are not observed in extremely premature neonatal aspirate samples

Previous reports have identified sex disparities in prenatal and perinatal lung dysfunction, disease outcomes, especially premature infants with very low birth weight [35, 36]. In our cohort, we sought to elucidate any sex-specific signatures of airway cellular development that may be detectable by evaluating both cell type composition and expression profile differences that might be attributable to sex differences. Generating pseudobulk expression profiles from each cell type for each individual subject, we performed principal component analysis (PCA) to assess whether the variance in our dataset could be explained by sex differences and did not find sex as significantly correlated with any of the first 20 PCs. Rather, we found the major components of variance (21.9% and 7.6% of the variance explained by PC1 and PC2, respectively) primarily separated epithelial cells (ciliated and basal) from myeloid cells (Fig. 2A). Similarly, when we evaluated the correlation between pseudobulk expression profiles of each cell type per individual with hierarchical clustering we found that clustering was driven primarily by cell type, with the first division again separating myeloid cells from epithelial and cycling cells (Fig. S6). Even within cell types, stratification by sex was not apparent. Next, we determined the proportion of each represented cell type per subject. The distribution across subjects (Fig. 2B) and sex (Fig. 2C) was broadly similar, although one subject (“NEA_20”) had a disproportionately large number of basal cells. Nonetheless, the pseudobulk transcriptional profile of NEA_20 cells clustered well with other samples in all previous analyses (e.g. NEA_20 basal cells were most correlated with basal cells from NEA_33, Spearman’s rho = 0.73), suggesting that the elevated proportion of basal cells in NEA_20 could be explained by variability in cell recovery rather than being representative of an aberrant state or poor quality sample.

### Premature neonate aspirates are enriched for intermediate myeloid states relative to adult airway samples

To better frame our understanding of the premature neonatal monocyte, intermediate, and macrophage cells with respect to differentiated lung tissues, we integrated our data with two published scRNA-seq datasets of adult lung tissue: Travaglini et al. 2020 [28] and the healthy subjects from Habermann et al. [37]. While these samples represent a different airway sampling strategy, they present the opportunity to evaluate our samples in the full spectrum of differentiated states. We first integrated our data with all cells from these two atlases and found that our samples clustered well with the expected cell types from the lungs (Fig. S7). To hone in on differences in the myeloid lineages, we next subset all data sets to just myeloid types of interest. Using the original cell annotations provided for each of the datasets, we subset the adult lung tissues and our premature neonatal data to the myeloid cells of interest (Monocytes, Classical Monocytes, OLR1+ Classical Monocytes, Nonclassical Monocytes, Intermediate Monocytes, and Macrophages). After randomly subsampling each adult dataset to contain the same number of cells as our dataset (5,161 cells from each sample, 15,483 cells in total), we re-clustered the integrated data (Fig. 3A). Supporting our initial findings, our neonatal intermediates formed a distinct cluster of cells from the monocyte and macrophage populations. When assessing the distribution of each dataset across each identified cell type, we confirmed that the majority of our premature neonatal cells are intermediate and monocyte populations while adult samples were primarily composed of macrophage and monocytes (Fig. 3B, Fig. S8). However, we noted an enrichment of our premature neonatal intermediate cells within two of the identified macrophage populations. 30.1% of the Intermediate I (VDAC1) cells (482/1597) clustered with adult macrophages (labeled as Mac (+Inter I) in Fig. 3B), suggesting a subset of our Intermediate I (VDAC1) cells are more transcriptionally related to macrophages than earlier intermediate cells. Similarly, 10.6% (107/1008) of the Intermediate II (MIR222HG) cells clustered with adult macrophages, suggesting that a subset of Intermediate II (MIR222HG) are also more differentiated. Upon investigating these two macrophage populations, we found differential expression of interstitial macrophage marker *CD68* for Mac (+Inter I) cells (Fig. 3C, adjusted p-value = 0.013) and alveolar macrophage marker *ITGAX* (the gene for CD11c) for the Mac (+Inter II) cells (Fig. 3D, adjusted p-value = 0.01; Table S5; [38]), implying that our intermediate myeloid cells are actively differentiating towards these endpoints.

### Trajectory analysis identifies distinct pathways for interstitial and alveolar macrophage differentiation

While the premature neonate endotracheal aspirate scRNA-seq profiles indicate a large composition of myeloid cells, many of these cells lacked expression profiles consistent with differentiated cell types. Previous flow cytometric analyses have indicated that interstitial macrophages (not alveolar macrophages) are present in the airways at birth, while alveolar macrophages appear to develop over the first several days of life [18], but these populations are not otherwise characterized in human populations to our knowledge. Even less is known about the state of myeloid cells during the canalicular/saccular transition in humans. In order to further define these intermediate cells, we subset our dataset to only the monocyte, intermediate, and macrophage clusters and performed trajectory analysis. Here we used the “Potential of Heat-diffusion for Affinity-based Transition Embedding” (PHATE) algorithm [39] to perform dimensionality reduction (Fig. 4A). With this analysis we noted two clear branches from monocytes either towards Intermediates II and III or towards macrophages. A number of cells bridging the divide were also observed between Intermediate I and Intermediates II and III (Cluster 4, Fig. S9). Therefore, we used Slingshot [40] to draw three trajectories: one from monocytes to the intermediate types II and III (we termed this trajectory “Alveolar” as this appears to be the path to alveolar macrophage precursors), another from monocytes to macrophages (which we termed “Interstitial” as the non-alveolar macrophage population), and a third between Intermediate I and Intermediates II and III (which we termed “Bridging”, as it represents a potential alternate route between interstitial macrophage precursors and alveolar macrophage precursors).

**Figure 4:**
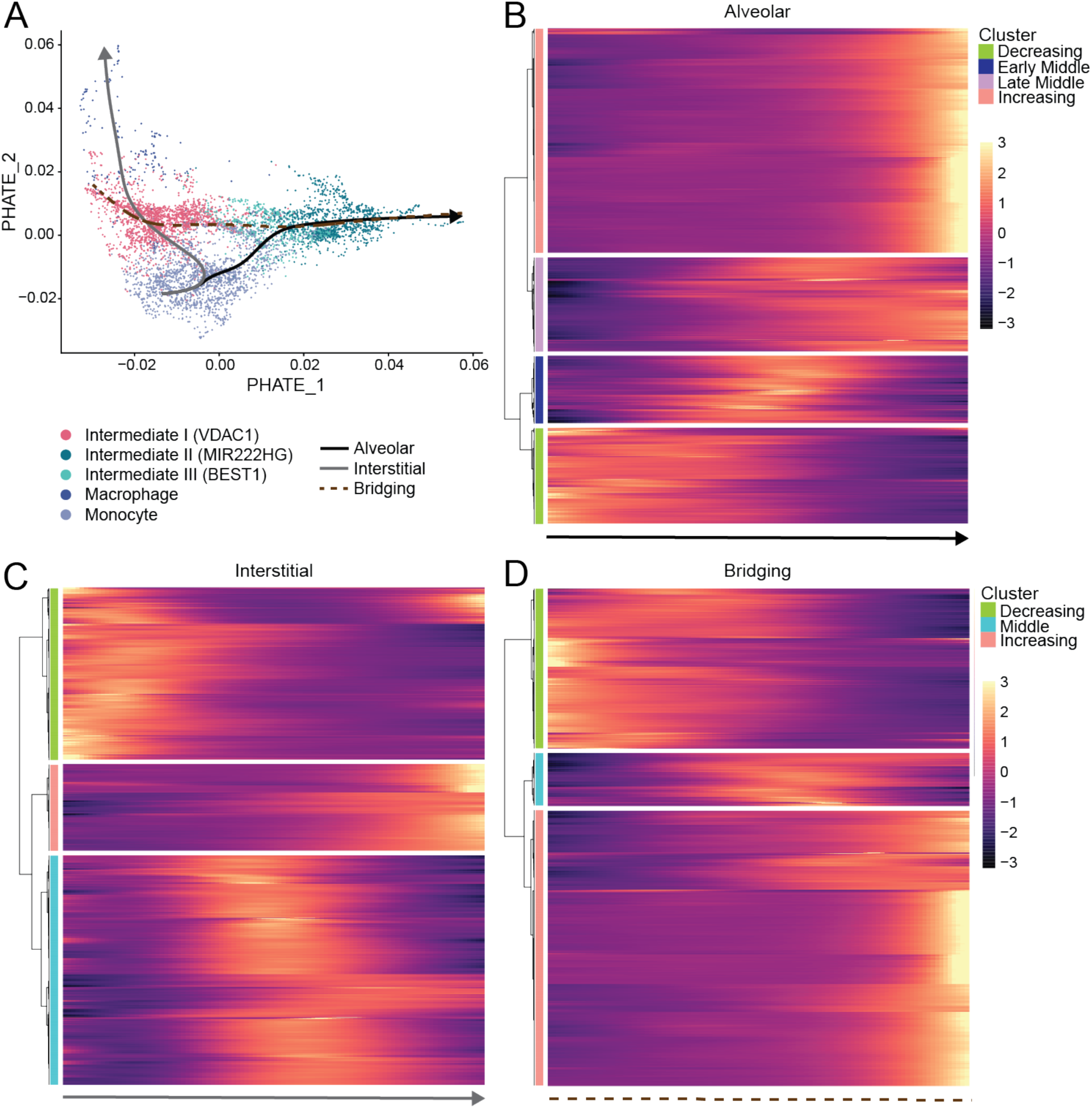
Trajectory analysis of premature neonate myeloid airway cells. (A) PHATE visualization of myeloid cells of interest colored by cell type annotation and visualization of Alveolar, Interstitial, and Bridging trajectory lineages. (B) Heatmap showing four major clusters of genes differentially expressed across Alveolar pseudotime. (C) Heatmap showing three major clusters of genes differentially expressed across Interstitial pseudotime. (D) Heatmap showing three major clusters of genes differentially expressed across Bridging pseudotime. For B-D, rows are individual genes, columns are points in pseudotime, and color ramp indicates scaled expression.

For the Alveolar trajectory, we identified 4,240 genes significantly differentially expressed (5% FDR) across pseudotime using Monocle 2 [41]. These genes exhibited 4 major expression patterns: 844 genes with decreasing expression across the trajectory (“Decreasing”), 590 genes with peak expression relatively early but in the middle of the trajectory but relatively early (“Early Middle”), 828 genes with peak expression in the middle of the trajectory but later than Early Middle genes (“Late Middle”), and 1,978 genes with increasing expression across pseudotime (“Increasing”) (Fig, 4B, Table S6). For each group of genes, we tested for enriched pathways (Table S7). Among the genes exhibiting a decreasing expression pattern across the trajectory we observed enrichments for ontology terms consistent with developing monocytes, such as “regulation of hematopoietic progenitor cell differentiation” (adjusted p-value = 4.0x10^-4^). Early Middle genes were enriched in pathways such as “cytokine-mediated signaling pathway” (adjusted p-value = 1.2x10^-9^) a process that is important for macrophage differentiation [42]. Genes with the Late Middle expression pattern showed enrichment for pathways such as “IRE1-mediated unfolded protein response” (adjusted p-value = 8.0x10^-3^), which is important in the process of polarization during macrophage maturation [43]. Genes with an Increasing expression pattern were enriched for pathways relevant to alveolar macrophages, including “Fc-gamma receptor signaling pathway involved in phagocytosis” (adjusted p-value = 7.3x10^-3^; [44]) and “regulation of cell-matrix adhesion” (adjusted p-value = 1.7x10^-2^; [45]).

Turning our attention to the Interstitial trajectory, we found a total of 3,696 genes to be significantly differentially expressed across pseudotime (5% FDR), comprising three major expression patterns: 1,302 “Decreasing” genes, 1,741 genes with the highest expression in the middle of the trajectory (“Middle”), and 653 “Increasing” genes (Fig. 4C, Table S6). As with the Alveolar trajectory, we tested for enriched pathways among the different classes of differentially expressed genes (Table S7). Decreasing genes were enriched for pathways including “positive regulation of interleukin-6 production” (adjusted p-value = 1.9x10^-10^), a pathway critical for development of monocytes into macrophages [46]. Analysis of Middle genes identified pathways including “ribosome biogenesis” (adjusted p-value = 7.8x10^-80^), which has been characterized as upregulated in “transitional” macrophages undergoing further terminal differentiation [47]. Increasing genes exhibited enrichments for “MHC class II antigen presentation” (adjusted p-value = 4.6x10^-6^), which is important for interstitial macrophage function [44].

To better understand the similarities and differences in expression patterns between these two trajectories, we intersected the genes differentially expressed along both trajectories. Due to the inherently conservative nature of significance testing, we used a two-threshold approach to be better powered to assess the overlap of these two trajectories (see Methods). Out of 6,113 genes that were differentially expressed along either trajectory at an FDR of 0.5, we identified 2,509 shared genes, 2,007 genes specific to the Alveolar trajectory, and 1,597 genes specific to the Interstitial trajectory (Table S6). Gene ontology analysis of genes specific to a given trajectory or shared (Table S8) revealed that the Alveolar trajectory-specific genes were enriched for “Chromatin organization” (adjusted p-value = 2.3x10^-8^) and “histone H4-K16 acetylation” (adjusted p-value = 3.6x10^-7^), consistent with studies showing that dysregulation of chromatin remodeling is implicated in experimental models and patients with hypoxia lung injury [48] and that H4K16ac regulates differentiation and apoptosis in embryonic myeloid cells [49]. Genes specific to the Interstitial trajectory remained enriched for pathways in relevant cellular metabolism terms like “mitochondrial translational elongation” (adjusted p-value = 2.2x10^-16^; [50]) and “rRNA processing” (adjusted p-value = 3.9x10^-7^; [51]). Enriched among genes shared between the Alveolar and Interstitial trajectories are pathways involved in lung inflammation response by recruitment of neutrophils such as “neutrophil activation involved in immune response” (adjusted p-value = 1.3x10^-52^) and “neutrophil mediated immunity” (adjusted p-value = 1.3x10^-51^), a well-characterized macrophage function [52]. Reassuringly, when we identified differentially expressed transcription factors (TFs) that were shared or unique (Table S6), we found several expected master regulators. For example, TFs specific to the Alveolar trajectory included *PPARG*, a TF with a well-characterized role in the differentiation of fetal monocytes into alveolar macrophages [53], as well as *KLF7*, which was also identified as an alveolar macrophage marker in a recent single-cell RNA-seq study [38]. TFs specific to the Interstitial trajectory included *SP1*, which has been shown to regulate *IL-10* production [54], a marker of interstitial macrophages [44]. Shared between the trajectories were TFs relevant to the differentiation of monocytes into macrophages such as *MEF2A* and *MEF2D* [55].

As the PHATE map indicated a potential secondary path linking the Alveolar and Interstitial trajectories, we also calculated pseudotime on this path (Methods) to draw the Bridging trajectory. We observed 4,566 genes significantly differentially expressed in three major classes: 1,495 genes that were upregulated in the Intermediate I population, 490 “Middle Bridging” genes, and 2,581 genes that were upregulated in the Intermediate II/III population (Fig. 4D, Table S6). Interestingly, among other pathways, Middle Bridging genes were enriched (Table S7) for pathways such as “VEGFA-VEGFR2 Signaling Pathway” (adjusted p-value = 1.2x10^-5^), consistent with the fact that alveolar macrophages produce *VEGF* during lung injury [56], and “regulation of TORC1 signaling” (adjusted p-value = 7.7x10^-3^), which has been reported as essential for alveolar macrophage development [57]. To get a better sense of what distinguishes this alternative trajectory linking alveolar and interstitial macrophages, we again employed our two-threshold approach to define shared and lineage specific transcriptional responses. Among the 6,661 genes differentially expressed in at least one trajectory, 285 were specific to the Bridging trajectory, while 259 were specific to the Alveolar trajectory, and 666 were specific to the Interstitial trajectory (Table S6). The remaining 5,451 were shared between at least two trajectories (1,246 were shared across all three trajectories), and were enriched (Table S8) for pathways such as “neutrophil degranulation” (adjusted p-value = 9.5x10^-43^), consistent with the enrichment of similar pathways among genes shared between Alveolar and Interstitial trajectories. We were particularly interested in the genes specifically differentially expressed in the middle of the Bridging trajectory, as these might highlight regulators of an alternative differentiation pathway between alveolar and interstitial macrophages. Of the 285 Bridging trajectory-specific genes, 101 exhibited peak expression in the cluster of cells bridging the Alveolar and Interstitial trajectories (Methods). While these genes were only moderately enriched (Table S8) for a single pathway (“Proteasome Degradation”, adjusted p-value = 0.04), we did note the presence of some genes with known roles in neonatal airway health (Table S6). One example was *MYC*, whose downstream targets have been shown to be upregulated in the macrophages of patients developing BPD [58], suggesting that this alternative pathway may play a role in BPD progression. In addition, some less characterized TFs that may play important roles (e.g. *ZNF672*, *ZFX*, and *ZGPAT*) were dynamic along this trajectory. In sum, our data reveal the initial differentiation steps that fetal monocytes have taken in the airways by the time of the canalicular/saccular transition in preparation for becoming mature alveolar macrophages after birth, and present three potential routes by which monocytes may become macrophages in premature neonatal airways.

## Discussion

In this study we generated single-cell profiles from endotracheal aspirates collected upon intubation of extremely premature neonates within the first hour of life. In spite of the challenges of working with these sample types (mucus, dead cells, sample availability logistics, etc.), we demonstrate that we are able to generate high quality data, although on relatively few cells per sample. With these data, we have developed a reference dataset of airway cells at birth in extremely premature neonates. Furthermore, by integrating our data with adult sample types, we are able to observe distinct subsets of intermediate myeloid cells in the developing lung preferentially clustering with two terminally differentiated macrophage populations from adults, further supporting our view of the differentiation trajectories of these cells. Using pseudotime analysis of our premature neonatal cells, we found one trajectory from monocytes to interstitial macrophages, one trajectory from monocytes to alveolar macrophage precursors, and one potential trajectory that may be an alternative differentiation pathway between alveolar and interstitial macrophage precursors. Interestingly, another recent single-cell study also proposed a secondary differentiation pathway between terminal macrophage fates that they observed across several tissues [59], suggesting that this may be a general feature of myeloid differentiation. While several previous studies have characterized myeloid populations from the airways in the days after birth [58, 60], to the best of our knowledge, this work represents the first transcriptional characterization of differentiating fetal monocytes in the human airway - the cells thought to give rise to alveolar macrophages in the first days after birth [61]. Further studies will be required to determine how these fetal human monocytes continue to differentiate throughout gestation and eventually give rise to alveolar macrophages after birth. One additional interesting implication of our results, consistent with other reports [62], is that monocytes appear to take up residence in the airspaces prior to differentiation (at least in part), suggesting that signaling in this compartment is necessary for myeloid differentiation, in contrast to lymphoid cells that are much less common in our analysis.

There are several important caveats to the work we have presented. First, we note that the viability for samples can vary widely. We collected samples from several patients that had even lower numbers of viable cells and so were not profiled for the purposes of this study. We believe that the source of many of the non-viable cells is the *in utero* swallowing of amniotic fluid, which contains dead cells from many epithelial sources. Whether this can be improved requires further experimentation. It is also important to acknowledge that the sample types presented here (newly generated endotracheal aspirates and published lung tissue samples) are not the only sampling method for surveying the airway lumen. Bronchoalveolar lavage [61, 62], as well as endobronchial biopsies and brushes [63] have all been used for single-cell characterization of the airways, although sampling the airways of extremely premature (or full-term) neonates is not trivial with any method. Additionally, the development of cells within our studies likely do not fully recapitulate normal airway development as these samples were collected from extremely premature neonates in respiratory distress. Nonetheless, given the challenges of sampling lungs at this critical time of development, we feel that these techniques (limited as they are) can offer an important view on biology. Furthermore, because samples were collected shortly after birth in premature subjects, fully differentiated alveolar macrophages are not expected in the airway [18]. In future studies, we hope to collect samples at later time points so that we can map the full trajectory of this lineage. In contrast to the existing literature [35, 36], we did not detect differences stratified by the sex of the subjects. We suspect that this might be due to the limited sample size in the current study. Future studies on larger populations will help to clarify this discrepancy. Also of note, some of our inferences were more difficult to corroborate - e.g., there is much more literature on alveolar macrophages than interstitial macrophages (and in some cases the two are not separated in studies, although they are readily identifiable as distinct clusters in many single-cell experiments [30,64,65]). There are also limitations to the directionality of our inferences. While our biological priors allowed us to place monocytes at the root of our Alveolar and Interstitial trajectories, the direction of differentiation is less clear for the Bridging trajectory. We attempted to implement an RNA velocity algorithm, but it failed to generate interpretable results (data not shown). Further work will be required to determine the cause of this failure. Finally, the comparison between the two sample types assessed in this study (adult lung tissue and premature neonate tracheal aspirates) is susceptible to confounding given the differences in protocol and the potential for differences in cell types or states that may be present in distinct sampling sites, and so these results should be viewed as preliminary. Nonetheless, as we highlighted, our results were largely consistent with the expected biology in terms of both the cellular composition and in age-related gene expression patterns, lending credence to the other differences observed.

Our broader goal is to extend the utility of single-cell assays into translational research tools and diagnostic development frameworks, and so we sought to develop a toolkit that would allow for single-cell profiling of the health of the airway from the moment of birth through adulthood, particularly in the event that patients may require intubation for respiratory distress. We hope that future studies will benefit from the application of these methods in specific disease settings to better understand disease pathogenesis and cellular states that might indicate future health problems or promising targets for novel interventions.

## Acknowledgements

We thank members of the Cusanovich and Ahmed labs. We thank B. Fransway, R. Sprissler, and J. Galina-Mehlman at the University of Arizona Genetics Core (UAGC) for assistance with sequencing. We thank N. Banovich and C. Trapnell for advice on the manuscript.

## Approvals

The University of Arizona Institutional Review Board approved the study and informed assent was obtained for all participants.

## Data and materials availability

Processed data will be made available through GEO upon publication. Raw data will be made available through dbGaP. Code will be made available at https://github.com/cusanovichlab/aspirates.

## Funding

HW was supported in part by an NHLBI fellowship (T32HL007249). Aspects of this work were supported by a University of Arizona Health Sciences Career Development Award (DAC).

## Supplementary Materials

### Supplementary Tables

**Table S1:** Endotracheal Aspirate Samples. This table is provided as a separate file. The columns are labeled by each subject ID (NEA_13, NEA_15, NEA_20, NEA_22, NEA_23, NEA_26, NEA_28, NEA_29, NEA_31, NEA_33). The rows are labeled as following: Row 1: Total Cell Count Upon Receipt Row 2: Estimated % Viability Upon Receipt Row 3: Mean Reads per Cell Row 4: Fraction of Reads in Cell Row 5: Recovered Cells Row 6: QC Filtered Cells Row 7: Recovered Median UMIs per Cell Row 8: QC Filtered Median UMIs per Cell Row 9: Recovered Median Genes per Cell Row 10: QC Filtered Median Genes per Cell Row 11: Median % MT Reads per Cell. Row 12: Subject Sex. Row 13: Sample Processing Batch.

**Table S2:** Cell Type Annotation and Label Transfer. Label transfer was performed in Seurat using Travaglini et al. [28] and the data from Adams et al. 2020 [29] as reference and using the R package SingleR [30]. For each cluster of cells, the most commonly assigned annotation was considered the overall annotation for that cluster (percent of cells annotated to the most common assignment provided in parentheses). Prediction scores of less than 0.5 for each cell were assigned to a “Low Score” category. If the majority of the cluster was assigned to the “Low Score” category, the second highest score has been included. After evaluating the results of both label transfers and examining relevant marker genes, a final label was determined for the cluster.

| Cluster | Travaglini et al. 2020 | Adams et al. 2020 | SingleR | Final Label |
| --- | --- | --- | --- | --- |
| 1 | OLR1+ Classical Monocyte (46.5%) | cMonocyte (52.6%) | Neutrophils (68.9%) | Neutrophil |
| 2 & 7 | Low Score (43.6%) / OLR1+ Classical Monocyte (30%) | Macrophage (81.9%) | Monocyte (89.5%) | Intermediate I (VDAC1) |
| 3 & 6 | Classical Monocyte (59.5%) | cMonocyte (74.7%) | Monocyte (89.5%) | Monocyte |
| 4 | Low Score (48.1%) / OLR1+ Classical Monocyte (27.1%) | Macrophage (85.1%) | Monocyte (73.7%) | Intermediate II (MIR222HG) |
| 5 | Low Score (40.4%) / OLR1+ Classical Monocyte (39.6%) | Macrophage (81%) | Monocyte (87.6%) | Intermediate III (BEST1) |
| 8 | Differentiating Basal (70.6%) | Low Score (48.9%) / Goblet (48.5%) | Epithelial_cells (98.7%) | Basal |
| 9 | Low Score (39.5%) / EREG+ Dendritic (31.6%) | Macrophage (63.8%) | Monocyte (42.4%) | Macrophage |
| 10 | Ciliated (80.5%) | Ciliated (88.7%) | Epithelial_cells (98.5%) | Ciliated |
| 11 | Myeloid Dendritic Type 2 (41.7%) | cDC2 (38.9%) | Monocyte (61.1%) | Cycling |

**Table S3:** Significant differentially expressed neonatal aspirate cell type markers. This table is provided as a separate file. The table includes the following columns: Column 1: “Gene” - differentially expressed gene name. Column 2: “AvgLogFC” - average log fold-change in gene expression level in the cell type specified in Column 5 compared to all other cell types. Column 3: “Pval” - unadjusted p-value. Column 4: “AdjPval” - Adjusted p-value using Benjamini-Hochberg method. Genes with an adjusted p-value < 0.05 are included in this table. Column 5: “CellType” - Cell type in which the differentially expressed gene in Column 1 was identified.

**Table S4:** Significant differentially expressed genes between neonatal monocyte and macrophages. This table is provided as a separate file. The table includes the following columns: Column 1: “Gene” - differentially expressed gene name. Column 2: “AvgLogFC” - average log fold-change in gene expression level between Monocytes and Macrophages (cell type defined in Column 5). Column 3: “Pval” - unadjusted p-value. Column 4: “AdjPval” - Adjusted p-value using Benjamini-Hochberg method. Genes with an adjusted p-value < 0.05 are included in this table. Column 5: “CellType” - Cell type for which the maker gene in Column 1 was identified.

**Table S5:** Significant differentially expressed genes between Mac (+Inter I) and Mac (+Inter II). This table is provided as a separate file. The table includes the following columns: Column 1: “Gene” - differentially expressed gene name. Column 2: “AvgLogFC” - average log fold-change in gene expression level between Mac (+Inter I) and Mac (+Inter II) (cell type defined in Column 5). Column 3: “Pval” - unadjusted p-value. Column 4: “AdjPval” - Adjusted p-value using Benjamini-Hochberg method. Genes with an adjusted p-value < 0.05 are included in this table. Column 5: “CellType” - Cell type for which the maker gene in Column 1 was identified.

**Table S6:** Pseudotime genes from the Alveolar trajectory, Interstitial trajectory, Bridging trajectory, two-way (Alveolar and Interstitial trajectories) and three-way (Alveolar, Interstitial, and Bridging trajectories) comparisons. This table is provided as a separate file. The columns of this table are as follows: Column 1: “Gene” - differentially expressed gene name. Column 2: “AM_Pval” - Alveolar trajectory unadjusted p-value. Column 3: “AM_Qval” - Alveolar trajectory q-value from Benjamini-Hochberg correction of p-values. Genes with a q-value < 0.05 are included in this table. Column 4: “AM_Cluster” - assigned Alveolar trajectory cluster for the heatmap in Fig. 4B. Column 5: “IM_Pval” - Interstitial trajectory unadjusted p-value. Column 6: “IM_Qval” - Interstitial trajectory q-value from Benjamini-Hochberg correction of p-values. Genes with a q-value < 0.05 are included in this table. Column 7: “IM_Cluster” - assigned Interstitial trajectory cluster for the heatmap in Fig. 4C. Column 8: “Bridging_Pval” - Bridging trajectory unadjusted p-value. Column 9: “Bridging_Qval” - Bridging trajectory q-value from Benjamini-Hochberg correction of p-values. Genes with a q-value < 0.05 are included in this table. Column 10: “Bridging_Cluster” - assigned Bridging trajectory cluster for the heatmap in Fig. 4D. Column 11: “TwoWay_Category” - category (AM_Specific, IM_Specific, Shared, or N/A) assigned to the gene in Column 1 when comparing the Alveolar and Interstitial trajectories. Column 12: “ThreeWay_Category” - category (AM_Specific, Bridge_Specific, IM_Specific, Shared, or N/A) assigned to the gene in Column 1 when comparing all three trajectories, Alveolar, Bridging, and Interstitial trajectories. Column 13: “GeneFullName” - full name of the gene in Column 1. Column 14: “GeneSynonym” - aliases of the gene in Column 1. Column 15: “TranscriptionFactor” - annotated “Yes” if the gene is a validated transcription factor. Column 16: “TF_PMID” - PubMed ID of scientific publications containing evidence of transcription factor validation. Column 17: “BridgingRegion” - annotated “Yes” if the gene maximum spline-estimated expression level was within the same window as the Cluster 4 pseudotime range. Column 18: “MaxSpline_Expression” - the maximum spline-estimated expression level of Bridging_Specific genes as specified in Column 12. Column 19: “MaxSpline_Bin” - the corresponding pseudotime bin value (1-100) of the maximum spline-estimated expression level in Column 18.

**Table S7:** Gene ontology terms from the Alveolar trajectory, Interstitial trajectory, and Bridging trajectory. This table is provided as a separate file. The columns of this table are as follows: Column 1: “Term” - gene ontology term. Column 2: “Database” - database which the term in Column 1 is defined. Column 3: “PathwayGenes” - total number of genes per term. Column 4: “AM_Cluster” - assigned Alveolar trajectory cluster for the heatmap in Fig. 4B. Column 5: “AM_Pval” - Alveolar trajectory unadjusted p-value for Fisher’s exact test or hypergeometric test. Column 6: “AM_Qval” - Alveolar trajectory q-value from Benjamini-Hochberg correction of p-values. Terms with a q-value < 0.05 are included in this table. Column 7: “AM_GeneHits” - total number of Alveolar genes identified for the pathway term. Column 8: “AM_Genes” - Alveolar gene names identified for the pathway term. Column 9: “AM_OddsRatio” - rank score or z-score for the deviation from an expected rank computed using a modification to Fisher’s exact test. Column 10: “AM_CombinedScore” - combined score by multiplying the natural logarithm of the p-value and z-score. Column 11: “IM_Cluster” - assigned Interstitial trajectory cluster for the heatmap in Fig. 4C. Column 12: “IM_Pval” - Interstitial trajectory unadjusted p-value for Fisher’s exact test or hypergeometric test. Column 13: “IM_Qval” - Interstitial trajectory q-value from Benjamini-Hochberg correction of p-values. Terms with a q-value < 0.05 are included in this table. Column 14: “IM_GeneHits” - total number of Interstitial genes identified for the pathway term. Column 15: “IM_Genes” - Interstitial gene names identified for the pathway term. Column 16: “IM_OddsRatio” - rank score or z-score for the deviation from an expected rank computed using a modification to Fisher’s exact test. Column 17: “IM_CombinedScore” - combined score by multiplying the natural logarithm of the p-value and z-score. Column 18: “Bridging_Cluster” - assigned Bridging trajectory cluster for the heatmap in Fig. 4D. Column 19: “Bridging_Pval” - Bridging trajectory unadjusted p-value for Fisher’s exact test or hypergeometric test. Column 20: “Bridging_Qval” - Bridging trajectory q-value from Benjamini-Hochberg correction of p-values. Terms with a q-value < 0.05 are included in this table. Column 21: “Bridging_GeneHits” - total number of Bridging genes identified for the pathway term. Column 22: “Bridging_Genes” - Bridging gene names identified for the pathway term. Column 23: “Bridging_OddsRatio” - rank score or z-score for the deviation from an expected rank computed using a modification to Fisher’s exact test. Column 24: “Bridging_CombinedScore” - combined score by multiplying the natural logarithm of the p-value and z-score.

**Table S8:** Pathway analysis of genes identified in two-way (Alveolar and Interstitial trajectories) and three-way (Alveolar, Interstitial, and Bridging trajectories) comparisons. Column 1: “Term” - gene ontology term. Column 2: “Database” - database which the term in Column 1 is defined. Column 3: “PathwayGenes” - total number of genes per term. Column 4: “TwoWay_Cluster” - category (AM_Specific, IM_Specific, Shared, or N/A) assigned to the term in Column 1 when comparing the Alveolar and Interstitial trajectories. Column 5: “TwoWay_Pval” - unadjusted p-value for Fisher’s exact test or hypergeometric test. Column 6: “TwoWay_Qval” - q-value from Benjamini-Hochberg correction of p-values. Terms with a q-value < 0.05 are included in this table. Column 7: “TwoWay_GeneHits” - total number of genes identified for the pathway term. Column 8: “TwoWay_Genes” - gene names identified for the pathway term. Column 9: “TwoWay_OddsRatio” - rank score or z-score for the deviation from an expected rank computed using a modification to Fisher’s exact test. Column 10: “TwoWay_CombinedScore” - combined score by multiplying the natural logarithm of the p-value and z-score. Column 11: “ThreeWay_Cluster” - category (IM_Specific, Shared, or N/A) assigned to the term in Column 1 when comparing all three trajectories, Alveolar, Bridging, and Interstitial trajectories. No terms were found for the Alveolar and Bridging specific genes in this analysis. Column 12: “ThreeWay_Pval” - unadjusted p-value for Fisher’s exact test or hypergeometric test. Column 13: “ThreeWay_Qval” - q-value from Benjamini-Hochberg correction of p-values. Terms with a q-value < 0.05 are included in this table. Column 14: “ThreeWay_GeneHits” - total number of genes identified for the pathway term. Column 15: “ThreeWay_Genes” - gene names identified for the pathway term. Column 16: “ThreeWay_OddsRatio” - rank score or z-score for the deviation from an expected rank computed using a modification to Fisher’s exact test. Column 17: “ThreeWay_CombinedScore” - combined score by multiplying the natural logarithm of the p-value and z-score. Column 18: “BridgingRegion_Cluster” - annotated “Yes” if the term in Column 1 specifies the pathway enriched in the Bridging Region (genes identified in Table S6, Column 17) Column 19: “BridgingRegion_Pval” - unadjusted p-value for Fisher’s exact test or hypergeometric test. Column 20: “BridgingRegion_Qval” - q-value from Benjamini-Hochberg correction of p-values. Terms with a q-value < 0.05 are included in this table. Column 21: “BridgingRegion_GeneHits” - total number of genes identified for the pathway term. Column 22: “BridgingRegion_Genes” - gene names identified for the pathway term. Column 23: “BridgingRegion_OddsRatio” - rank score or z-score for the deviation from an expected rank computed using a modification to Fisher’s exact test. Column 24: “BridgingRegion_CombinedScore” - combined score by multiplying the natural logarithm of the p-value and z-score.

### Supplementary Figures

**Figure S1:**
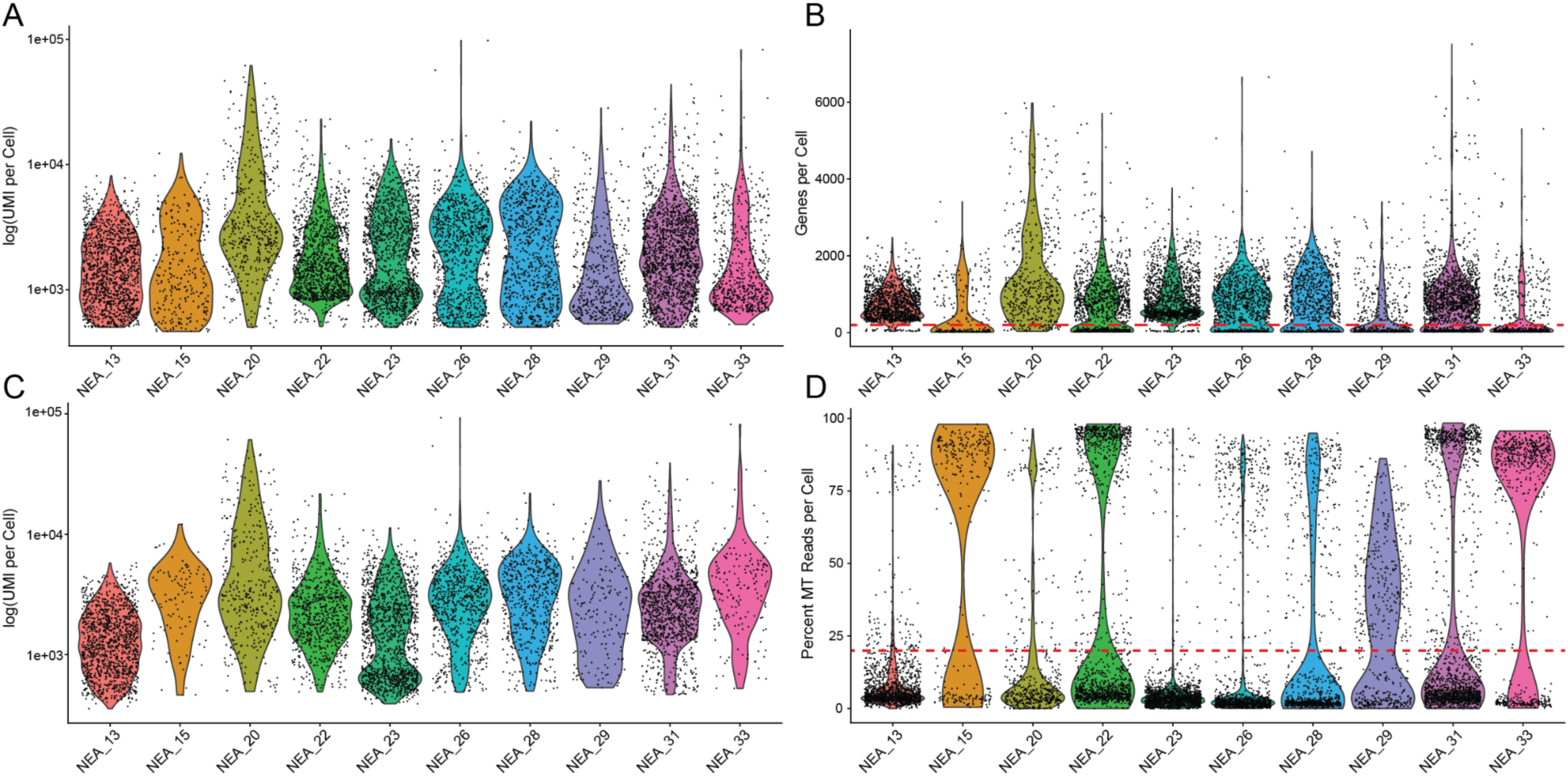
Single-cell data quality per subject and quality filtering. (A) Violin plot of the unfiltered, log-scaled Unique Molecular Identifiers (UMIs) per recovered cell by each subject, with each point representing an individual cell. (B) Violin plot of the recovered genes per cell separated by subject. Cells with less than 200 genes per cell (read dashed line) were quality filtered. (C) Violin plot of the log-scaled UMIs per cell across subjects after quality filtering. (D) Violin plot of the percent mitochondrial (MT) reads per cell by each subject. Cells with greater than 20% (red dashed line) were removed from downstream analysis.

**Figure S2:**
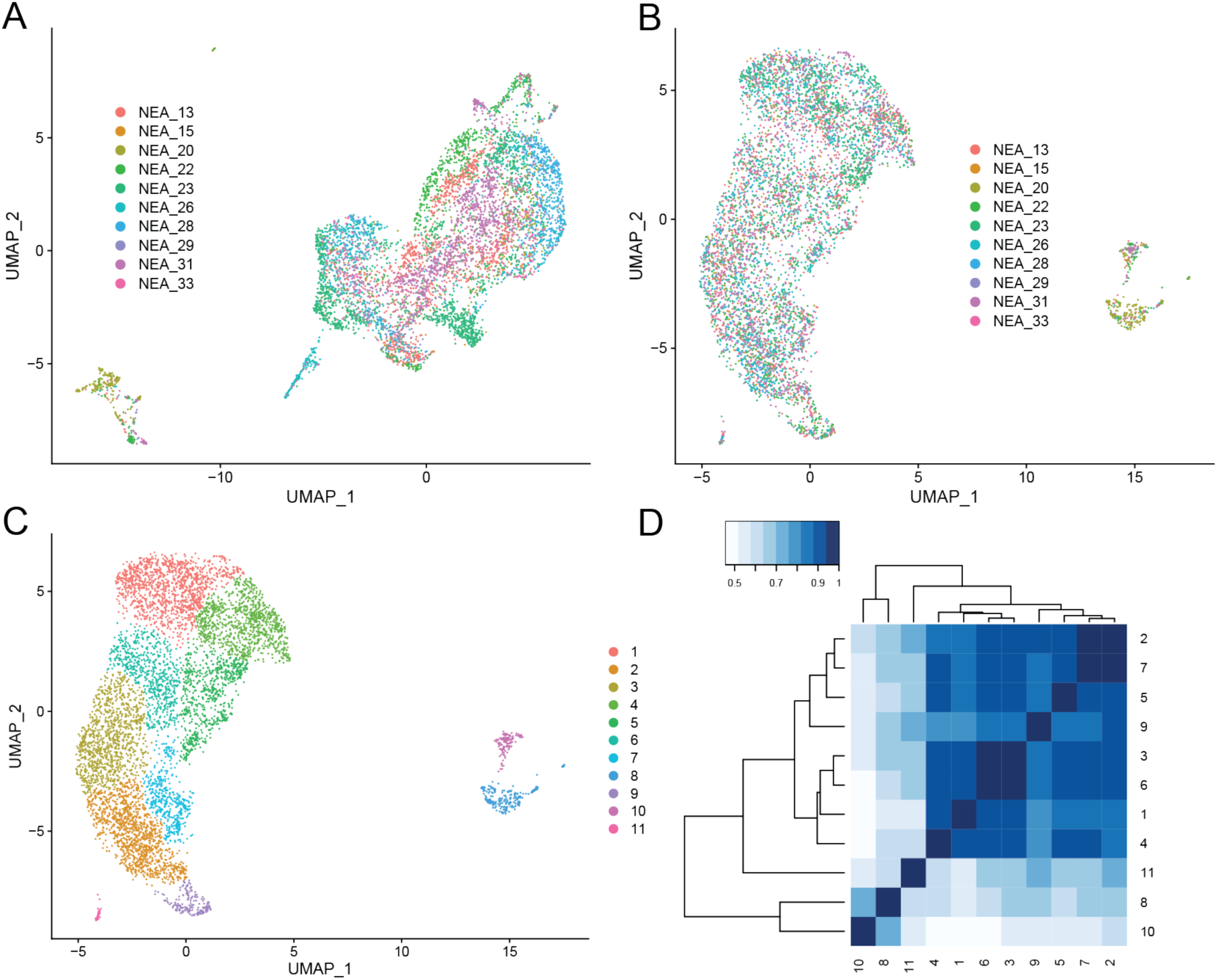
Initial clustering, Harmony integration, and cluster correlation. (A) UMAP representation of subjects without Harmony integration. (B) UMAP reduction after Harmony integration labeled by subject. (C) UMAP representation of the 11 original identified clusters. (D) Heat map of Spearman correlation between identified clusters.

**Figure S3:**
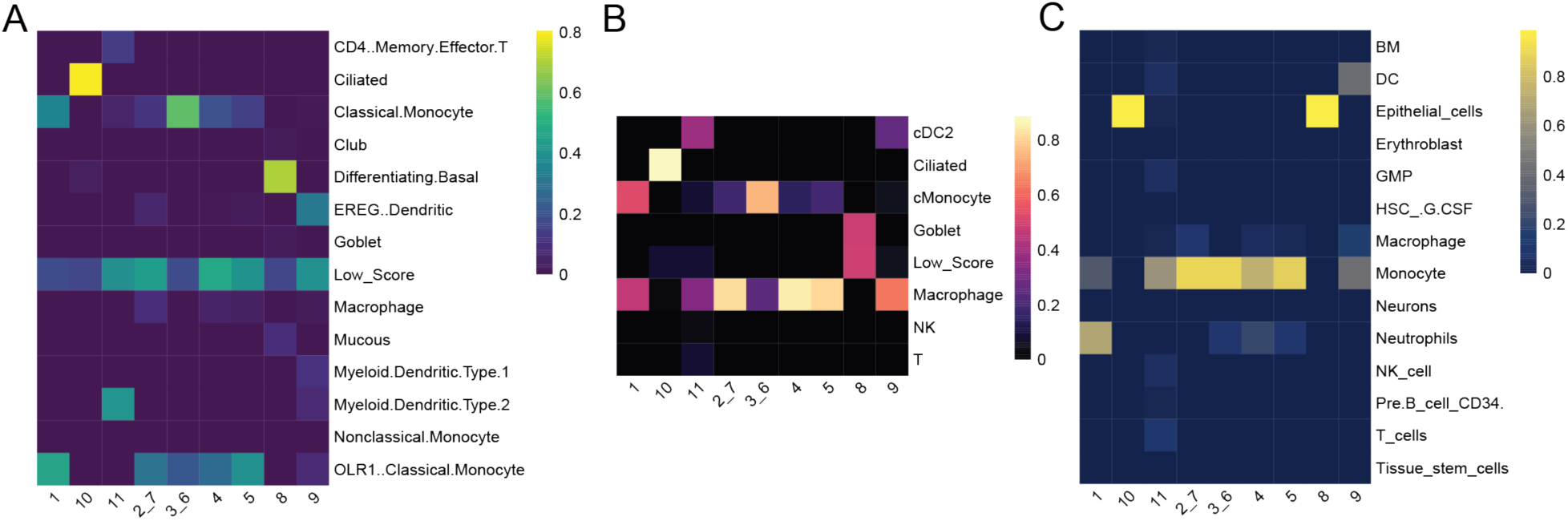
Neonatal label transfer and cell type assignment. (A) Confusion matrix of the Travaglini et al. cell type annotation transfer. Rows represent clusters identified in data. Columns represent cell types defined in Travaglini et al.. Color scale indicates proportion of cells within a cluster assigned to a given annotation. (B) Confusion matrix of the cell type labels transferred from Adams et al. 2020 (C) Confusion matrix of the SingleR cell type annotation transfer.

**Figure S4:**
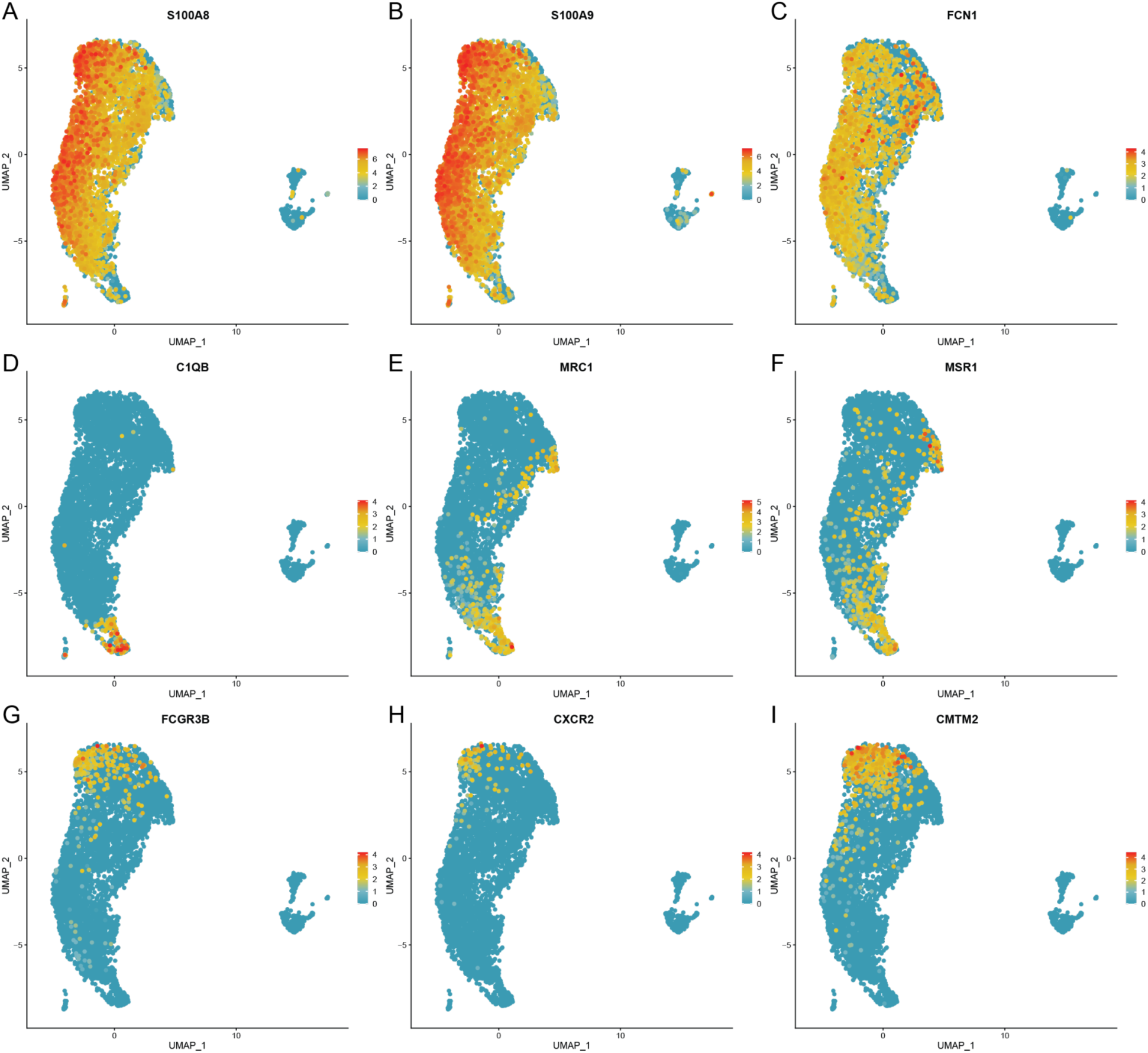
Myeloid markers across myeloid cell types. (A-C) UMAP colored by expression level for monocyte marker genes *S100A8* (A), *S100A9* (B), and *FCN1* (C). (D-F) UMAP colored by expression level for differentiated cell marker genes *C1QB* (D), *MRC1* (E), and *MSR1* (F). (G-I) UMAP colored by expression level for neutrophil marker genes *FCGR3B* (G), *CXCR2* (H), and *CMTM2* (I).

**Figure S5:**
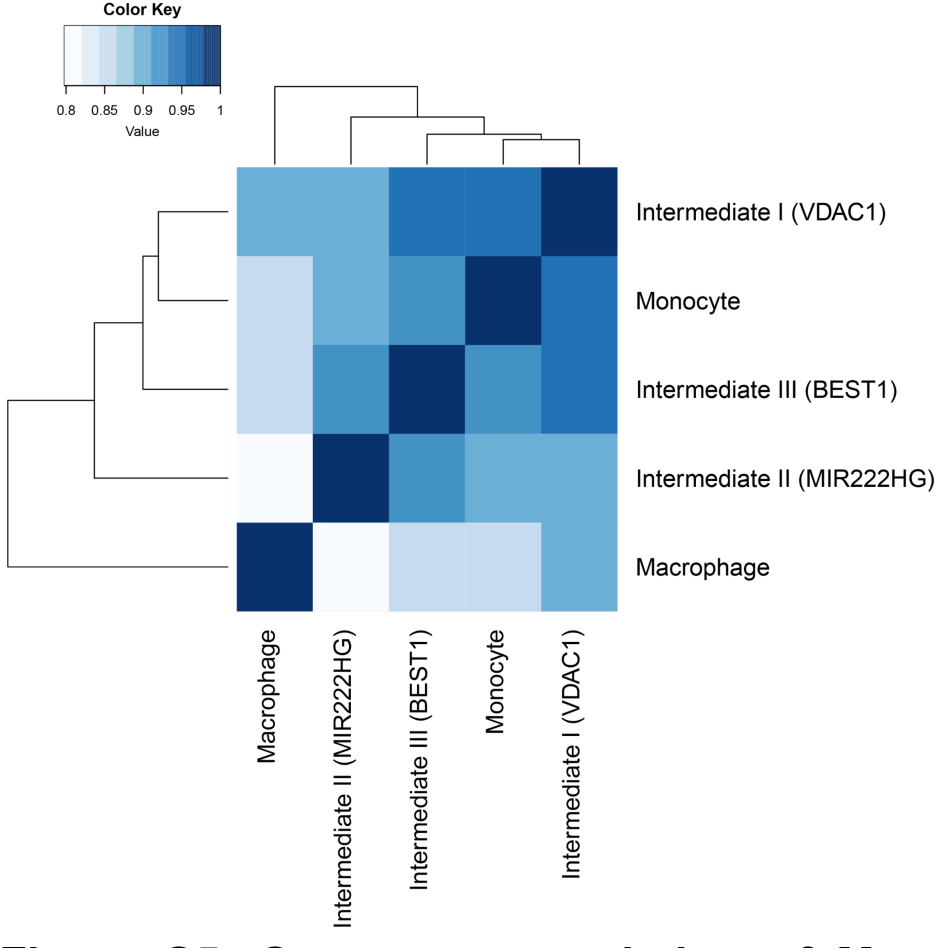
Spearman correlation of Monocytes, Intermediates, and Macrophages. (A) Heatmap of the Spearman correlation between Monocytes, Macrophages, Intermediate I (VDAC1), Intermediate II (MIR222HG), and Intermediate III (BEST1).

**Figure S6:**
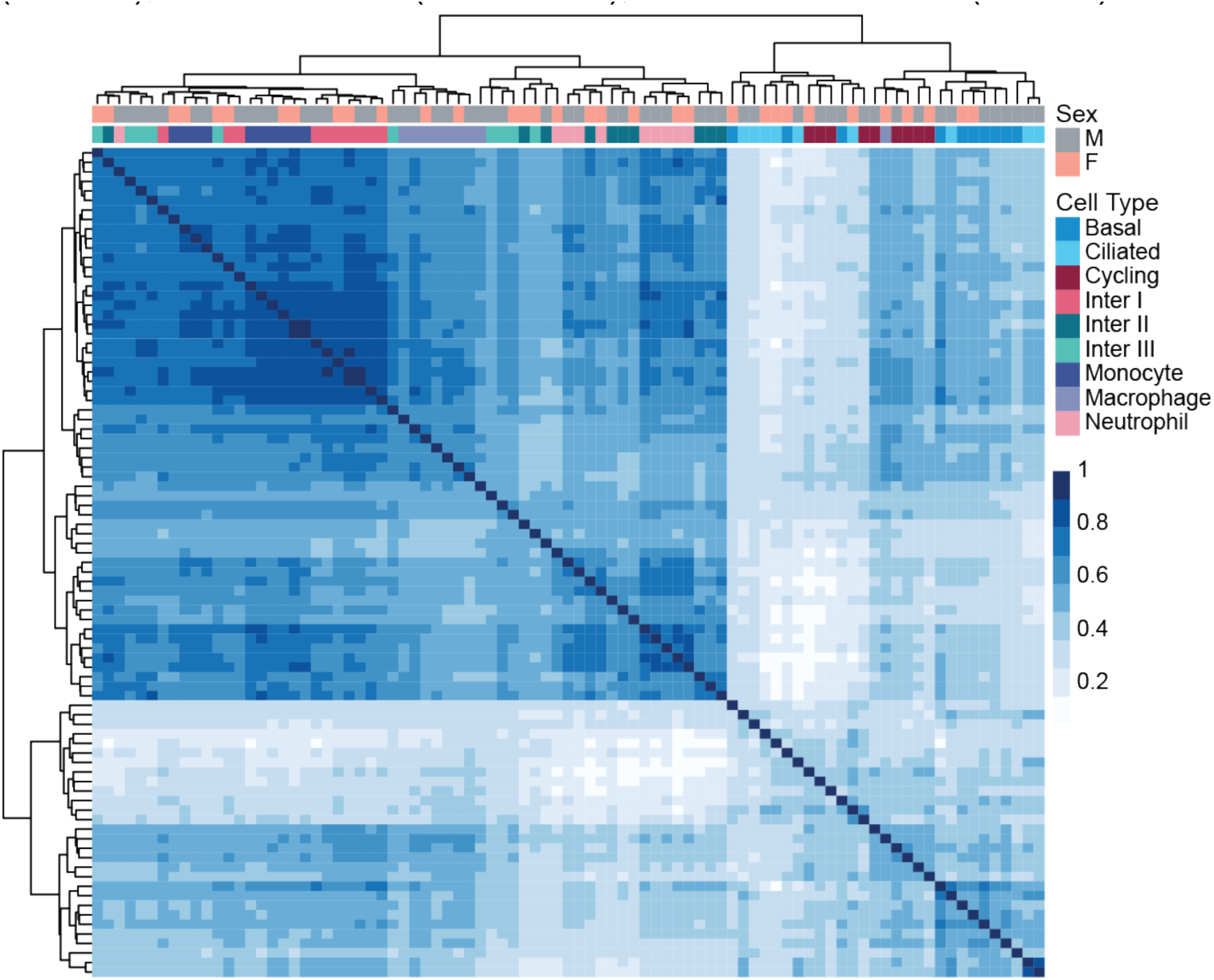
Patient cell type and sex correlation. Spearman correlation matrix between cell types by individual pseudobulk profiles. Colored bars represent sex of subject and cell type.

**Figure S7:**
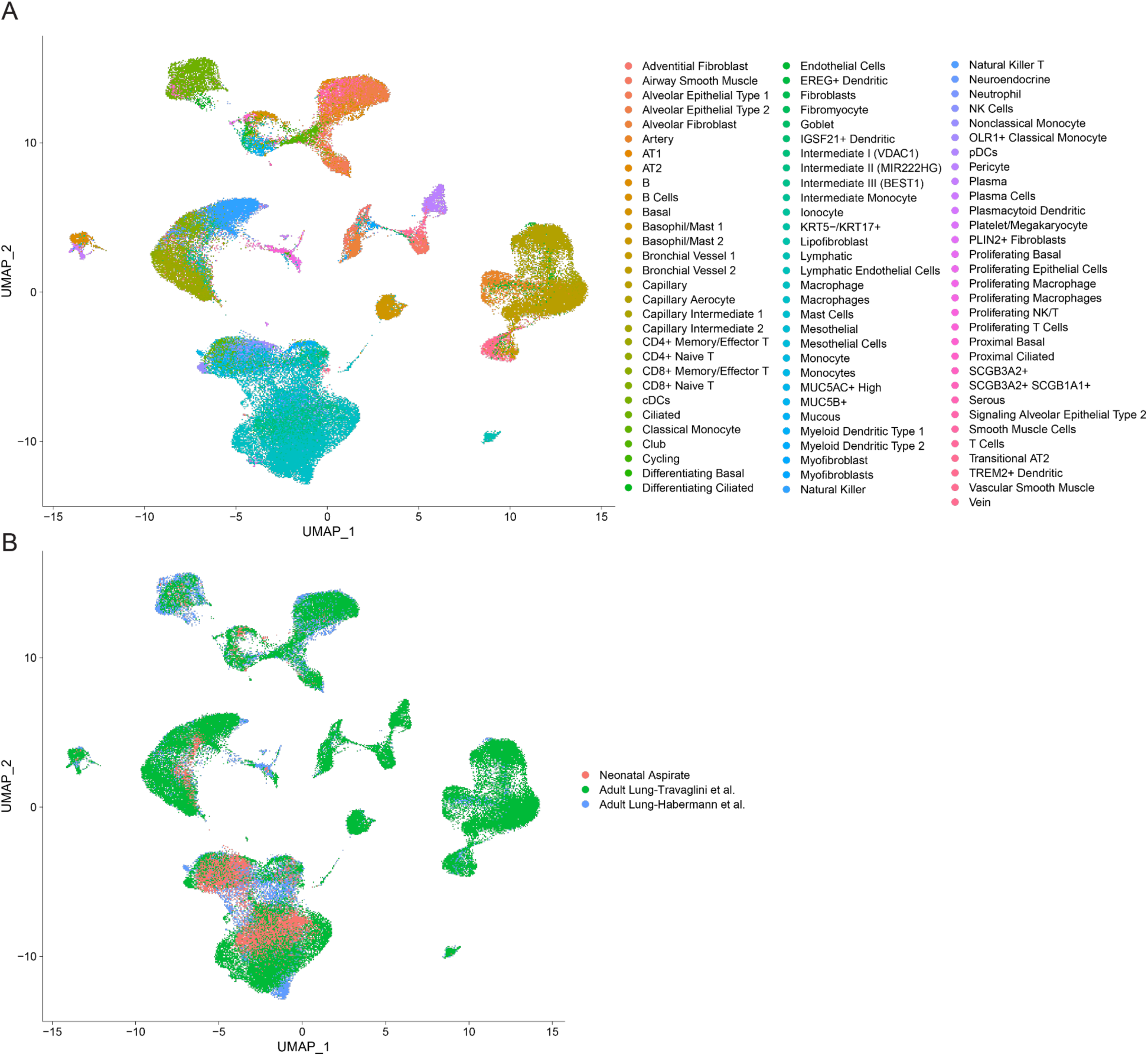
Integration of neonatal aspirate, Travaglini et al., and Habermann et al. datasets. (A) UMAP representation of integrated neonate and adult data colored by the original cell type annotations from each study. (B) UMAP representation of integrated neonate and adult data colored by the original dataset.

**Figure S8:**
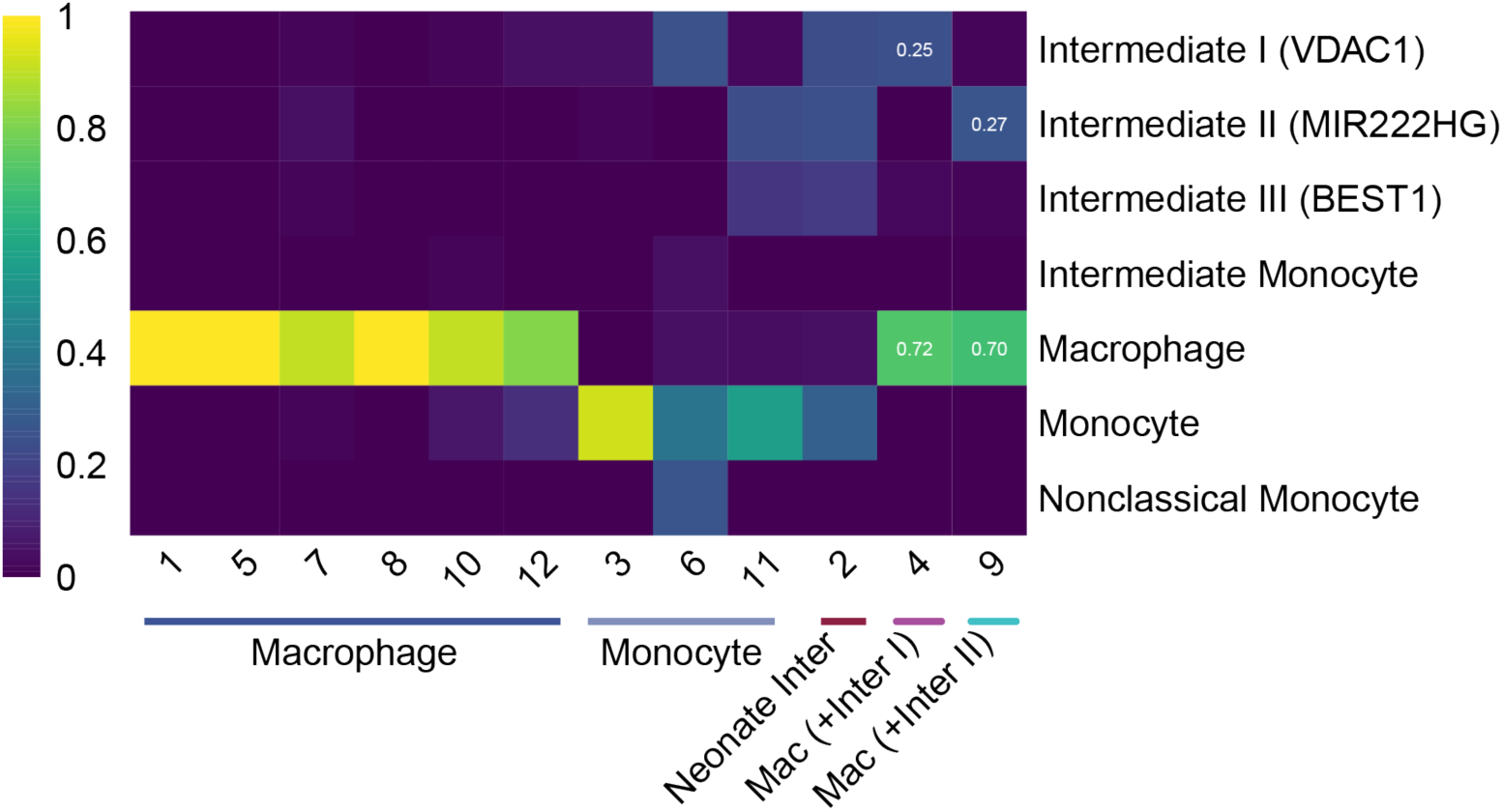
Distribution of neonatal aspirate, Travaglini et al., and Habermann et al. cells across myeloid clusters. Confusion matrix showing the proportion of each original cell type annotation (rows) by each cluster (columns). Final cluster cell type annotation provided below cluster value. Values highlighted for Mac (+Inter I) and Mac (+Inter II).

**Figure S9:**
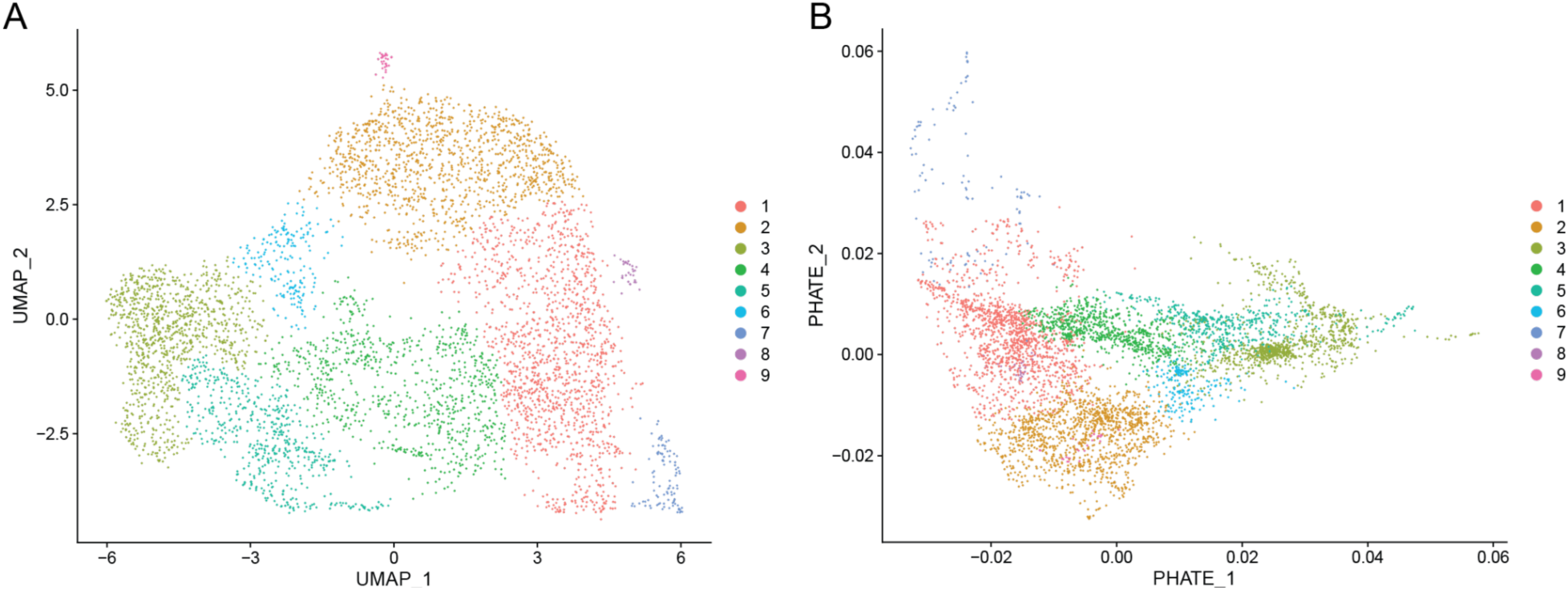
Neonatal monocyte, intermediate, and macrophage cells. (A) UMAP representation showing the clusters identified in neonatal Monocytes, Intermediate I (VDAC1), Intermediate II (MIR222HG), Intermediate III (BEST1), and Macrophages. (B) PHATE reduction colored by clusters annotated in (A). Cluster 4 represents the intermediate Bridging cells.

## Methods

### Endotracheal Aspirate Collection

Endotracheal aspirates were collected from premature neonates (23-28 weeks gestational age, gestational birth weight < 1250 gram) in whom endotracheal tubes had been placed for clinical reasons. Within 60 minutes of birth, while still in the delivery suite, aspirate samples were collected immediately following placement of the tracheal tube before administering any surfactants or other medications. The aspirate samples were collected by instilling a 1 mL saline solution into the tracheal tube followed by immediate suction. After 1-2 mL of DMEM (Corning 10-013-CV) was added to the sample, it was stored at 4°C until it could be transferred on ice to the laboratory for cryopreservation.

### Endotracheal Aspirate Processing

After receipt from the clinic, each sample was transferred to a 50mL conical tube and a viability count was estimated with Trypan Blue staining using the Countess II FL. Samples were washed with 10 volumes of ice-cold DPBS and were centrifuged at 500 x g for 5 minutes at 4°C, aspirated, and the cell pellet was resuspended in 1 mL freezing media (RPMI 40% FBS 10% DMSO). Mucus in the samples was relatively minor. Samples were transferred to a cryotube and frozen overnight in CoolCells (Corning) in the -80°C freezer, and then transferred to liquid nitrogen the following day for storage. On the day of library generation, samples were thawed quickly in a 37°C water bath, transferred to 10mL media (RPMI 15% FBS), and centrifuged at 500 x g for 5 minutes at 4°C. After, the cells were resuspended in 0.04% BSA in PBS, counted on a Countess (Bio-Rad) after staining with Trypan Blue, and adjusted to a concentration of 1x10^6^ cells/mL.

### 10X Library Generation

All libraries were generated using the 10X Chromium Single Cell 5ʹ GEM, Library & Gel Bead Kit v1.1 across two batches with five samples in each batch. Based on the cell counts described above, the maximum number of cells (16,000) were loaded for each sample, unless the concentration was too low, in which case 37.8 ul of the sample (the maximum volume possible in this reaction) was loaded regardless of concentration. Consequently, in the first batch, we loaded an estimated 16,000 cells for NEA_13, 16,000 cells for NEA_15, 4,914 cells for NEA_20, 4,158 cells for NEA_22, and 7,938 cells for NEA_23 on the Chromium Controller. In the second batch (processed at a later date), we loaded an estimated 12,474 cells for NEA_26, 10,584 cells for NEA_28, 11,340 cells for NEA_29, 16,000 cells for NEA_31, and 7,560 cells for NEA_33. All samples were processed according to the 10X protocol.

### Sequencing of 10X Libraries

Sequencing was performed in two different runs. Within each batch, samples were pooled at molar ratios proportional to the estimated number of cells loaded on the 10X lane for each sample. Batch 1 (NEA_13, NEA_15, NEA_20, NEA_22, and NEA_23) along with samples that did not ultimately meet our QC thresholds (NEA_09, NEA_10, NEA_19, and NEA_24) were sequenced using a NextSeq Mid Output Kit run with Read 1=26, Read 2=91, and Index 1=8. Batch 2 (NEA_26, NEA_28, NEA_29, NEA_31, and NEA_33) along with two samples that did not ultimately pass our QC thresholds (NEA_32 and NEA_34) were sequenced using a NextSeq High Output Kit (multiplexed with other samples not relevant to this study) with the following parameters: Read 1=51, Read 2=91, Index 1=10.

### Analysis of Sequencing Data

#### Preprocessing Single-cell RNA-seq Data

Raw sequencing data were processed using the 10X Genomics Cell Ranger pipeline. FASTQ files for Batch 1 were generated with default arguments. FASTQ files of Batch 2 samples were generated with Cell Ranger ‘mkfastq’ and the base mask argument ‘--use-bases-mask’ to create FASTQ files of Read 1=26 and Index 1=8. Preliminary quality control metrics (fraction of reads in cells, reads per cell, etc.) were collected from the Cell Ranger Web Summary. Sparse matrices of UMI counts for each gene observed in cells were generated and subsequently analyzed with other programs. To adjust for cell-free RNA contamination, the Cell Ranger outputs were loaded into the R package SoupX [66] using the ‘load10x’ function. Ambient RNA contamination was estimated with ‘autoEstCont’ function and argument tfidfMin = 0.7 for all samples and decontaminated with the function ‘adjustCounts.’

#### Analysis in Seurat

The contamination-adjusted matrices generated through the 10X Cell Ranger pipeline and SoupX were loaded into the R package Seurat v3 [67]. Samples were removed from the study if they did not meet the following criteria: a minimum fraction of reads in cells of 0.4 (related to ambient RNA contamination) per sample and a minimum of 100 total cells per sample after filtering out cells with less than 200 genes per cell and a mitochondrial read fraction less than 0.2 (indicative of dead or dying cells [66]). Doublets were identified using the Python package Scrublet v0.2.1 [68]. Each sample was analyzed for potential doublets using the default pipeline as described in the Scrublet GitHub repository applied to the 10X Cell Ranger output matrix (i.e. prior to SoupX correction). The cell barcodes identified as potential doublets were filtered out of each Seurat object. The Seurat objects were then merged and normalized using the ‘NormalizeData’ function. We then estimated the top 3,000 variable genes using the ‘FindVariableFeatures’ function and the argument selection.method = "vst". Subsequently, the data were scaled using the ‘ScaleData’ function and PCA was run using the ‘RunPCA’ function using the variable genes identified by ‘FindVariableFeatures’. Samples were integrated using Harmony [69] with the ‘RunHarmony’ wrapper function in Seurat. The Harmony integrated components were used as input for UMAP dimensionality reduction with the first 20 dimensions using the ‘RunUMAP’ function. The K-nearest neighbors of each cell were found and used to construct a Shared Nearest Neighbor graph under the function ‘FindNeighbors’ with the first 2 dimensions. To identify clusters, we used the ‘FindCluster’ function using the Leiden algorithm with the resolution 0.12 for the UMAP in Fig. 1A. The sex of subjects was inferred from sequencing data by assessing the expression level of Y chromosome genes *DDX3Y*, *KDM5D*, *ZFY*, and *EIF1AY*, all of which were consistently expressed across 6 subjects.

#### Label Transfer and Cell Type Annotation

We then performed label transfer using two reference datasets following the Seurat vignette and the R package SingleR [30]. The first reference dataset was from the Travaglini et al. [28] and the counts, cell IDs, and metadata were downloaded from Synapse (syn21041850). The data were used to generate a Seurat object (data were normalized, variable features were found, data were scaled, PCA was run, neighbors and clusters were found, and UMAP was run). Using this as our reference dataset, we found label transfer anchors with the first 30 dimensions using the ‘FindTransferAnchors’ function in Seurat. These anchors were used in the ‘TransferData’ function to add predictions to our data from the cell type annotations stored in the metadata of the reference dataset. We determined the proportion of cells in each cluster assigned to each label and considered the most common label as the annotated cell type for clusters identified in our dataset. Cells with a prediction value of <0.5 were assigned as “Low Score” cells. A second label transfer was used for additional comparison from the scRNA-seq of healthy patients described in [29] - data downloaded from Gene Expression Omnibus (GEO) (GSE136831). First, a Seurat object was made from the counts matrix, cell IDs, and metadata and the healthy patient data were used by subsetting on the “Control” patients from the metadata table. Label transfer with Seurat was performed exactly as described for the Travaglini et al. dataset. For the final label transfer method, SingleR, we used the Human Primary Cell Atlas [70] reference data and used the default parameters for the ‘SingleR’ function. The majority cell type assignment for each cluster was assigned to the entire cluster. We used a combination of the three annotation results and differentially expressed marker genes in our clusters to make a final cell type annotation.

#### Integration with Adult Lung Datasets

We generated Seurat objects of the data from Travaglini et al., as described above, and data from the healthy controls in Habermann et al. Adult data were normalized independently and then merged with our neonatal data. Using the original cellular annotations of each dataset, we subset the data to the myeloid cells of interest. Each adult dataset was randomly subsampled to the same number of cells per dataset (5,161 cells from each study, 15,483 total). We estimated the top 2,000 variable genes, data were scaled, and PCA was run as outlined in “Analysis in Seurat.” Samples were integrated using Harmony, including dataset and individual as terms, with the first 20 PCs. UMAP was run, neighbors and clusters found using the Leiden algorithm with the resolution 0.45.

#### Differential Expression Analysis

Differentially expressed genes were identified using the ‘Libra’ R package [31]. We used the Libra function ‘run_de’ with our processed gene by cell matrix and appropriate metadata of cell type and replicate information to implement the edgeR likelihood-ratio test framework. Significant differential expression was defined as an adjusted p-value < 0.05.

#### Trajectory Analysis

For our trajectory analysis, we subset the data on the monocytes, macrophages, and intermediates from our neonatal data. UMAP was run and clusters were found with the resolution 0.09. We then used the R package PHATE [39] and the wrapper function ‘RunPHATE’ for dimensionality reduction. For Fig. 4, the Alveolar and Interstitial trajectories were calculated on this representation of the data excluding Cluster 4 (cells spanning between Intermediate I and Intermediate II and III). The Bridging trajectory was calculated on the PHATE reduction excluding Monocytes and Macrophages. Here we used the ‘slingshot’ function from the R package Slingshot [40] and the ‘start.clus’ parameter as Monocytes and ‘end.clus’ as Intermediate II (MIR222HG) and Macrophage for the Alveolar and Interstitial trajectories, and ‘start.clus’ as Intermediate I (VDAC1) and ‘end.clus’ as Intermediate II (MIR222HG) for the Bridging Trajectory. The pseudotime value matrix was extracted with the function ‘slingPseudotime’ and added as metadata in creating a Monocle CellDataSet object. Differentially expressed genes across pseudotime were found using Monocle 2. Briefly, the function differentialGeneTest was used with the Subject ID included as a covariate. Genes with an adjusted p-value < 0.05 were considered significant and were included in the heatmap in Fig. 4B-D (generated with the function ‘plot_pseudotime_heatmap’ and the number of clusters set to 4 in Fig. 4B and 3 in Fig. 4C-D).

#### Gene Ontology Enrichment

Significantly differentially expressed genes (defined above) were used as gene lists for gene ontology enrichment analysis. The enrichment tests were performed using the R package enrichR [71, 72] and testing for enrichment of annotations in the “GO_Biological_Process_2021”, "WikiPathway_2021_Human", and "Reactome_2016" databases. Results were filtered on an adjusted p-value < 0.05 for each ontology term and requiring 2 or more genes in a category.

#### Transcription Factor Identification

TFs within a list of genes were identified by using TFCheckpoint [73]. Only genes with experimental evidence for TF activity (regulation of RNA polymerase II and specific DNA binding activity) were considered as TF hits.

#### Differential Expression Sharing Across Trajectories

In order to define differentially expressed genes that were shared between trajectories, we used a two-threshold approach similar in spirit to the method used in [74] to define shared factor binding in a comparative ChIP-seq study. To implement this approach, we took all genes that were differentially expressed along a trajectory and then asked if they would have been considered differentially expressed along a second trajectory at a relaxed threshold (unadjusted p-value < 0.05). After performing the reciprocal intersection, all genes that overlapped in either direction were considered shared, while significant genes that were not identified at a relaxed threshold in the other trajectories were considered specific to a particular trajectory.

#### Differential Expression Across Cells Spanning the Bridging Trajectory

To define Bridging-specific genes that are most relevant to the cells spanning between Intermediate I and Intermediates II and III, we first determined the range of pseudotime values assigned to this cluster of cells (referred to as “Cluster 4” in *Trajectory Analysis* above) along the Bridging trajectory. We then subset the 285 genes identified as Bridging-specific such that only genes whose maximum spline-estimated expression level was within the same window as the Cluster 4 pseudotime range (bins 19 to 62) were considered relevant to the bridging region. Subsetting this way resulted in the identification of 101 genes that were specific to the Bridging trajectory and whose maximum expression was localized to the region spanning the Interstitial and Alveolar trajectories.

## Notes

### Competing Interest Statement

The authors have declared no competing interest.

